# Translation-independent association of mRNAs encoding protomers of the 5-HT_2A_-mGlu2 receptor complex in living cells

**DOI:** 10.1101/2024.06.17.599432

**Authors:** Somdatta Saha, Javier Gonzalez-Maeso

## Abstract

The serotonin 2A receptor (5-HT_2A_R) and the metabotropic glutamate 2 receptor (mGluR2) form heteromeric G protein-coupled receptor (GPCR) complexes through a direct physical interaction. Co-translational association of mRNAs encoding subunits of heteromeric ion channels has been reported, but whether complex assembly of GPCRs occurs during translation remains unknown. Our *in vitro* data reveal evidence of co-translational modulation in *5-HT_2A_R* and *mGluR2* mRNAs following siRNA-mediated knockdown. Interestingly, immunoprecipitation of either 5-HT_2A_R or mGluR2, using an antibody targeting epitope tags at their N-terminus, results in detection of both transcripts associated with ribonucleoprotein complexes containing RPS24. Additionally, we demonstrate that the mRNA transcripts of *5-HT_2A_R* and *mGluR2* associate autonomously of their respective encoded proteins. Validation of this translation-independent association is extended *ex vivo* using mouse frontal cortex samples. Together, these findings provide mechanistic insights into the co-translational assembly of GPCR heteromeric complexes, unraveling regulatory processes governing protein-protein interactions and complex formation.

## INTRODUCTION

The majority of proteins across all domains of life function as part of multimeric complexes (Goodsell and Olson, 2000; Marsh and Teichmann, 2015). G protein-coupled receptors (GPCRs) represent a highly diverse group of transmembrane proteins that exert various physiological effects (Pierce et al., 2002; Rosenbaum et al., 2009). This superfamily of membrane proteins can be grouped into six classes (A-F) based on their primary sequence and signaling properties (Gonzalez-Maeso and Sealfon, 2010; Pincas et al., 2017). Molecular and functional studies indicate that class C GPCRs such as GABA_B_ receptors and metabotropic glutamate receptors (mGluRs) behave as obligate dimers (Niswender and Conn, 2010; Pin and Bettler, 2016; Frangaj and Fan, 2018). Additionally, while class A GPCRs were traditionally recognized for their efficient functioning as monomers; at least when reconstituted into a phospholipid bilayer (Whorton et al., 2007; Whorton et al., 2008; Kuszak et al., 2009), there is a growing body of evidence suggesting the potential for class A GPCRs to form homodimers/heterodimers and oligomers. These complexes exhibit distinct pharmacological, trafficking, and functional properties compared to their parent monomeric forms (Milligan, 2009; Gonzalez-Maeso, 2011; Gaitonde and Gonzalez-Maeso, 2017; Ferre et al., 2022). Although the physiological relevance of GPCR heteromerization is not fully established, it holds promise for generating novel drug target combinations and fine-tuning the structure and function of one or more GPCRs involved in the complex to enhance therapeutic strategies.

The serotonin (5-hydroxytryptamine, or 5-HT) 2A receptor (5-HT_2A_R) which belongs to class A, along with the class C mGluR2 are GPCRs involved in processes related to cognition, perception, and mood regulation. They constitute significant targets for various psychoactive substances, including psychedelics (also known as classical hallucinogens) such as psilocybin or lysergic acid diethylamide (LSD), as well as antipsychotics such as clozapine or risperidone (Moreno et al., 2011; Moreno and Gonzalez-Maeso, 2013; Moreno et al., 2013; Gonzalez-Maeso, 2017; Shah and Gonzalez-Maeso, 2019; Jaster and Gonzalez-Maeso, 2023; Saha and Gonzalez-Maeso, 2023). Several lines of evidence suggest that these two neurotransmitter GPCRs can physically interact; influencing G protein coupling, function and trafficking (Gonzalez-Maeso et al., 2008; Fribourg et al., 2011; Moreno et al., 2012; Moreno et al., 2016; Shah et al., 2020). Insight into the role of this 5-HT_2A_R-mGluR2 complex in agonist-induced endocytic processes has been provided (Toneatti et al., 2020), yet much remains unknown about the assembly of their component protomers in terms of location, timing, and mechanism. Understanding this is crucial, as protein complexes serve as fundamental organizational units in the proteome, and their assembly within the crowded cellular environment is a complex task (Schwarz and Beck, 2019).

Cells employ various strategies to ensure faithful and efficient assembly, ranging from random subunit collisions (Hardy et al., 1988; Gingras et al., 2007) to the involvement of cellular chaperones (Doring et al., 2017; Rousseau and Bertolotti, 2018) and dedicated assembly organelles (Klinge and Woolford, 2019). Another mechanism involves immediate co-translational folding and the simultaneous association of binding partners, preventing premature or unintended interactions of nascent peptides. While early biochemical evidence in prokaryotes demonstrated co-translational enzymatic activity in nascent multimeric enzymes (Koubek et al., 2021), this concept has been extended to eukaryotic heteromeric complexes (Lin et al., 2000; Shiber et al., 2018; Badonyi and Marsh, 2022; Seidel et al., 2022). For example, it was established that alternative mRNA transcripts encoding *hERG1a* and *hERG1b* subunits, which assemble to produce the cardiac K_v_ channel (Sanguinetti and Jurkiewicz, 1990), are physically associated during translation (Liu et al., 2016). However, up to our knowledge, there is currently no biological evidence supporting the co-translational association of nascent GPCRs in mammalian cells.

Previous findings indicated a crosstalk mechanism between *5-HT_2A_R* and *mGluR2* affecting transcriptional processes. For instance, 5-HT_2A_R knockout mice exhibited reduced cortical expression of *mGluR2* mRNA (Kurita et al., 2013), and chronic treatment with atypical antipsychotics down-regulated cortical *mGluR2* mRNA via 5-HT_2A_R (Kurita et al., 2012; de la Fuente Revenga et al., 2019). Building on these observations, here we explored the possibility of co-translational association of mRNA encoding protomers of the heteromeric 5-HT_2A_R-mGluR2 complex.

## RESULTS

### Cross-regulation of mGluR2 mRNA and protein levels by 5-HT_2A_R

To investigate the reciprocal regulation of 5-HT_2A_R and mGluR2 at both the mRNA and protein levels, we aimed to elucidate the impact of individually silencing these genes in a co-expression system. Initially, we assessed the influence of *5-HT_2A_R* small interfering RNA (siRNA) and *mGluR2* siRNA in HEK293 cells transfected to co-express cMyc-5HT_2A_R and HA-mGluR2, respectively. Each siRNA, as compared to a non-targeting siRNA control, led to an approximately 50% reduction in the corresponding mRNA (Figure 1A, and Figure S1A), and immunoreactive levels (Figures 1B, 1C, and Figures S1B, S1C). Interestingly, silencing *5-HT_2A_R* with its specific siRNA in cells co-expressing 5-HT_2A_R and mGluR2 resulted in a significant decrease in *mGluR2* mRNA (Figure 1D) and immunoreactive levels (Figures 1E, 1F), each reduced by almost half. However, downregulation of *mGluR2* by its specific siRNA in cells co-expressing both receptors did not significantly impact the mRNA (Figure S1D) and immunoreactive levels (Figures S1E, S1F) of 5-HT_2A_R. To confirm the specificity of this cross-regulation between 5-HT_2A_R and mGluR2, we examined the expression of mGluR3 (mRNA and protein) upon downregulation of *5-HT_2A_R* in cells co-expressing cMyc-5HT_2A_R and HA-mGluR3. The expression of *mGluR3* mRNA (Figure 1G) and immunoreactivity (Figures 1H, 1I) remained unchanged following *5-HT_2A_R* silencing. Thus, the simultaneous decrease in *mGluR2* mRNA upon silencing of *5-HT_2A_R* mRNA suggests a potential physical association between the co-expressed transcripts. The concurrent reduction in their protein levels upon downregulation of 5-HT_2A_R further supports the idea of transcriptional regulation during translation of the interacting proteins.

**Figure 1.**
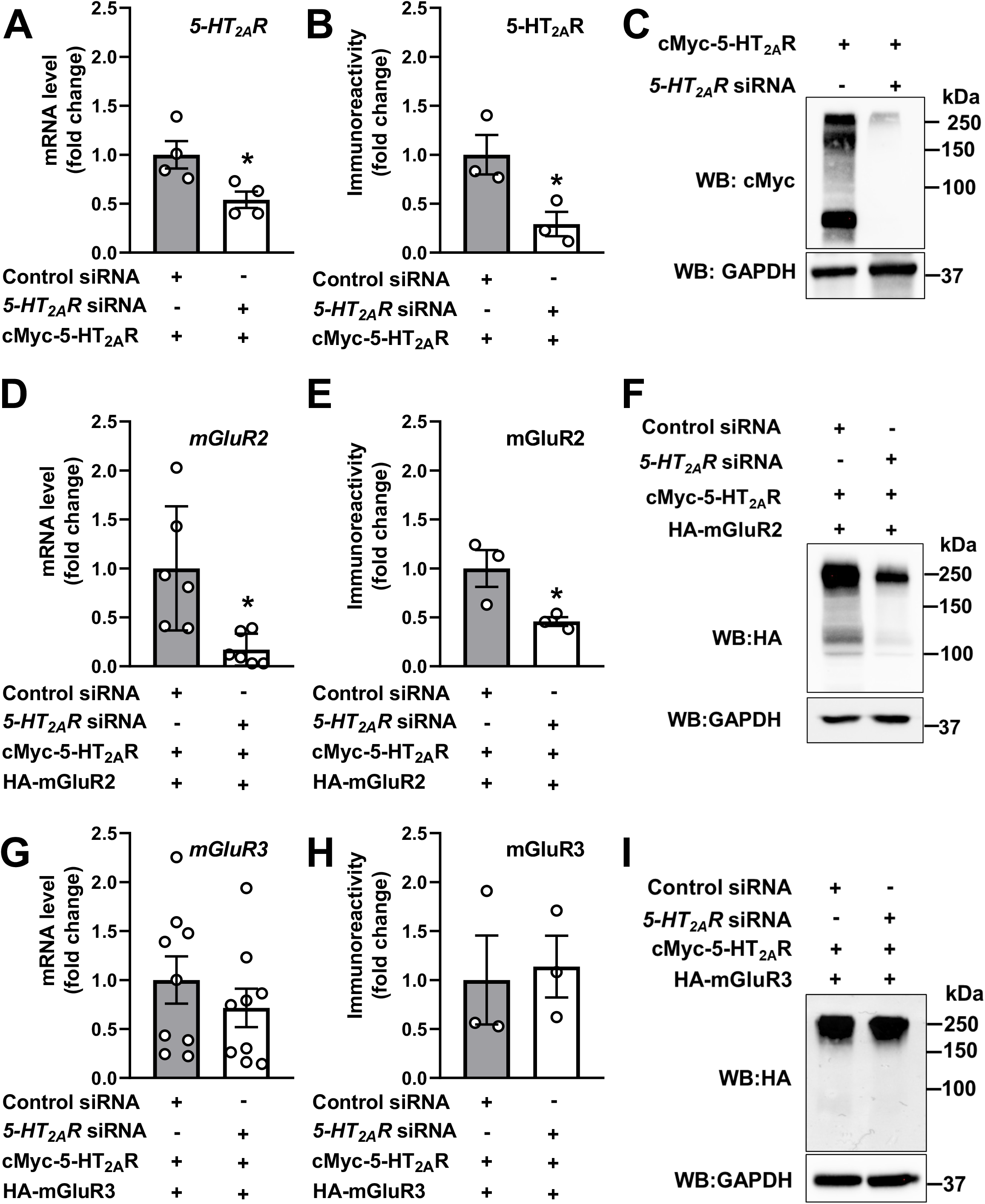
Cross-regulation of mGluR2 expression by 5-HT_2A_R. (**A-C**) HEK293 cells were transfected with non-targeting siRNA or *5-HT_2A_R* siRNA. Forty-eight hours after siRNA transfection, cells were transfected with pcDNA3.1-cMyc-5-HT_2A_R. RNA and protein extractions were carried out 24 h following DNA transfection. *5-HT_2A_R* mRNA was assessed by RT-qPCR (n = 4 independent experiments) (**A**), and 5-HT_2A_R immunoreactivity was assessed by Western blot (n = 3 independent experiments) (**B**). Representative immunoblots are shown (**C**). (**D-F**) HEK293 cells were transfected with non-targeting siRNA or *5-HT_2A_R* siRNA. Forty-eight hours after siRNA transfection, cells were co-transfected with pcDNA3.1-cMyc-5-HT_2A_R and HA-mGluR2. RNA and protein extractions were carried out 24 h following DNA transfection. *mGluR2* mRNA was assessed by RT-qPCR (n = 6 independent experiments) (**D**), and mGluR2 immunoreactivity was assessed by Western blot (n = 3 independent experiments) (**E**). Representative immunoblots are shown (**F**). (**G-I**) HEK293 cells were transfected with non-targeting siRNA or *5-HT_2A_R* siRNA. Forty-eight hours after siRNA transfection, cells were co-transfected with pcDNA3.1-cMyc-5-HT_2A_R and HA-mGluR3. RNA and protein extractions were carried out 24 h following DNA transfection. *mGluR3* mRNA was assessed by RT-qPCR (n = 9 independent experiments) (**G**), and mGluR3 immunoreactivity was assessed by Western blot (n = 3 independent experiments) (**H**). Representative immunoblots are shown (**I**). Unpaired two-tailed Student’s *t*-test (*p < 0.05).

### Co-translational assembly of *5-HT_2A_R* and *mGluR2* transcripts

We posited that if *5-HT_2A_R* and *mGluR2* transcripts physically associated during translation, it would be feasible to immunoprecipitate both transcripts using an antibody against the N-terminus of either of their nascent proteins. To assess this, HEK293 cells underwent co-transfection with cMyc-5-HT_2A_R and HA-mGluR2. Subsequently, ribonucleoprotein (RNP) complexes were extracted using a conventional RNP immunoprecipitation (RIP) assay, employing an anti-HA antibody for the immunoprecipitation of the nascent HA-mGluR2 polypeptide (for validation of the cellular fractionation protocol using markers of cellular compartments, including cytoplasmic *α*-Tubulin and nuclear Lamin A/C, see Figures S2A, S2B). This was followed by RNA isolation and reverse transcription PCR (RT-PCR) assays to identify transcripts associated with the mGluR2 protein (Figure 2A). Likewise, an anti-cMyc antibody was used to immunoprecipitate the nascent cMyc-5-HT_2A_R protein, and RNA isolation and RT-PCR assays were performed to identify transcripts linked with the 5-HT_2A_R protein (Figure 2B) (for immunoblots showing expression of the used constructs at the expected molecular weights, see Figures S2C, S2D).

**Figure 2.**
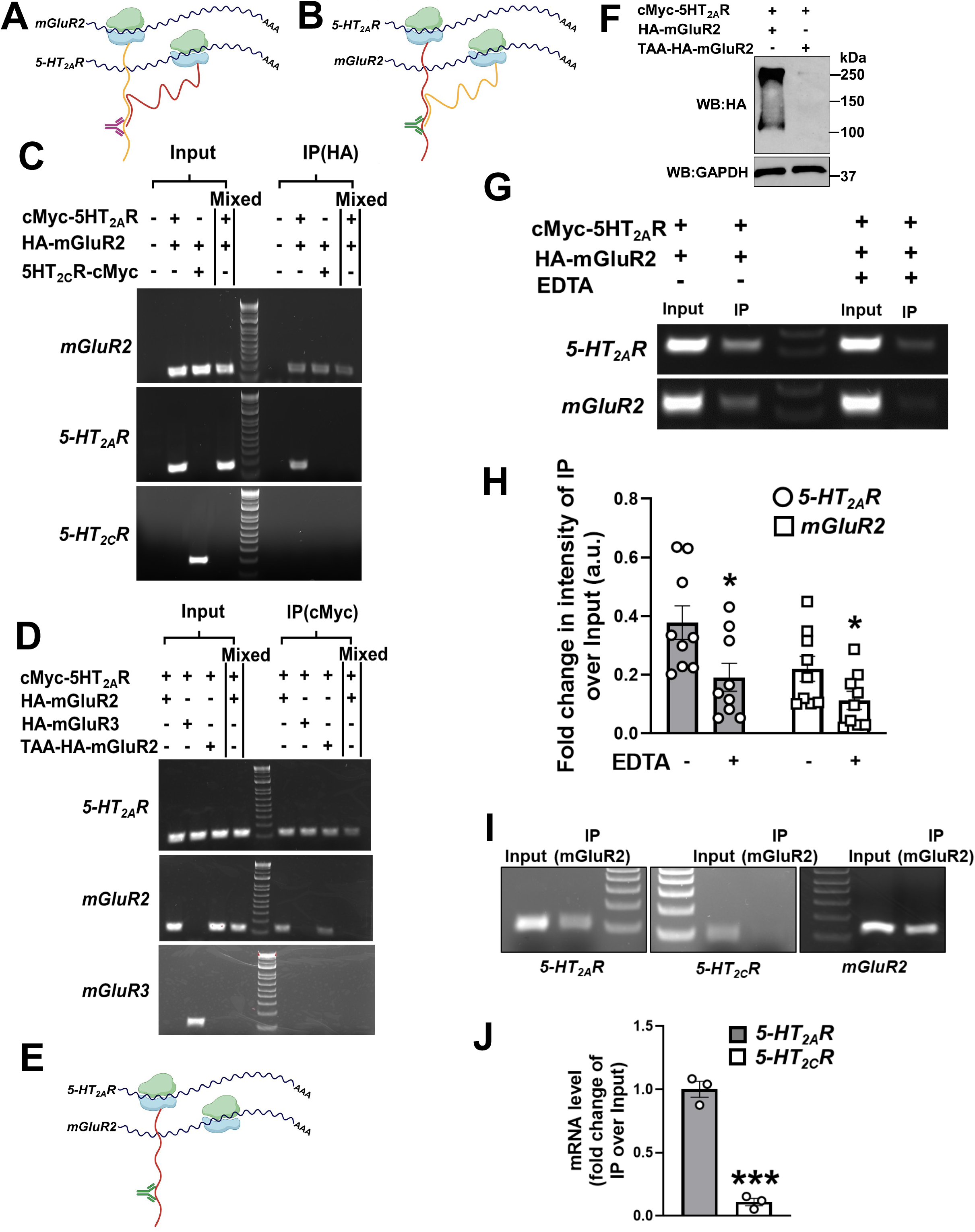
Co-translational assembly of *5-HT_2A_R* and *mGluR2* mRNAs. (**A**) Schematic representation illustrating the co-translational association of 5-HT_2A_R and mGluR2 polypeptides, depicted in red and yellow, respectively, as they emerge from ribosomes shown in blue and green. The polypeptides originate from neighboring *5-HT_2A_R* and *mGluR2* transcripts, depicted in black. The mGluR2 targeting antibody (anti-HA, magenta) used for immunoprecipitation of the mRNA/protein complex is indicated, bound to the N-terminal of mGluR2. (**B**) Schematic representation illustrating the co-translational association of 5-HT_2A_R and mGluR2 polypeptides, depicted in red and yellow, respectively, as they emerge from ribosomes shown in blue and green. The polypeptides originate from neighboring *5-HT_2A_R* and *mGluR2* transcripts, depicted in black. The 5-HT_2A_R targeting antibody (anti-cMyc, dark green) used for immunoprecipitation of the mRNA/protein complex is indicated, bound to the N-terminal of 5-HT_2A_R. (**C**) HEK293 cells were co-transfected with pcDNA3.1-HA-mGluR2, and pcDNA3.1-cMyc-5-HT_2A_R or pcDNA3.1-5-HT_2C_R-cMyc constructs, or mock. Images show representative RT-PCR products for *mGluR2*, *5-HT_2A_R* and *5-HT_2C_R* transcripts from HEK293 cells before (Input) and after immuno-precipitation (IP) using an anti-HA antibody. For control, cells separately expressing the c-Myc- or HA-tagged forms were mixed. Data are representative from three independent experiments. (**D**) HEK293 cells were co-transfected with pcDNA3.1-cMyc-5-HT_2A_R, and pcDNA3.1-HA-mGluR2, pcDNA3.1-HA-mGluR3 or pcDNA3.1-TAA-HA-mGluR2 constructs. Images show representative RT-PCR products for *5-HT_2A_R*, *mGluR2* and *mGluR3* transcripts from HEK293 cells before (Input) and after immuno-precipitation (IP) using an anti-cMyc antibody. For control, cells separately expressing the c-Myc- or HA-tagged forms were mixed. Data are representative from three independent experiments. (**E**) Schematic showing anti-cMyc antibody used to immunoprecipitate 5-HT_2A_R protein and association of *5-HT_2A_R* and *mGluR2* transcripts in the absence of mGluR2 protein. (**F**) Immunoblot demonstrating loss of HA-mGluR2 protein when the translation initiation site is mutated (TAA). **(G, H)** HEK293 cells were co-transfected with pcDNA3.1-cMyc-5-HT_2A_R and pcDNA3.1-HA-mGluR2 constructs. Images show representative RT-PCR products for *5-HT_2A_R* and *mGluR2* transcripts from HEK293 cells before (Input) and after immuno-precipitation (IP) using an anti-HA antibody. RIP assays were performed in presence and absence of EDTA (25 mM). (**H**) Quantification of change in IP band intensities of RT-PCR products for *5-HT_2A_R* and *mGluR2* transcripts as observed in presence and absence of 25mM EDTA. Representative image (**G**), and quantification of IP/input band intensities (n = 3 independent experiments performed in triplicate) (**H**). (**I**) Images show representative RT-PCR products for *5-HT_2A_R*, *5-HT_2C_R* and *mGluR2* transcripts from mouse frontal cortex samples before (Input) and after immuno-precipitation (IP) using an anti-mGluR2 antibody. Data are representative from three independent experiments in 3 mice. (**J**) Mouse frontal cortex samples were subjected to RIP assays employing an anti-mGluR2 antibody. Subsequently, the RNP complexes underwent processing for RNA isolation and RT-qPCR assays for the detection of *5-HT_2A_R* and *5-HT_2C_R* transcripts. Data are shown as fold change of IP/Input (n = 3 mice). Two-way ANOVA followed by Bonferroni’s *post-hoc* test (**H**), and unpaired two-tailed Student’s *t*-test (**J**). (*p < 0.05, **p < 0.001).

Notably, our findings indicate that the anti-HA antibody immunoprecipitates not just the mGluR2 protein but also the corresponding mRNAs for *mGluR2* and *5-HT_2A_R* (Figure 2C). Likewise, the anti-cMyc antibody is capable of immunoprecipitating not only the 5-HT_2A_R protein but also the associated *5-HT_2A_R* and *mGluR2* mRNAs (Figure 2D). Importantly, this association was not observed in cells co-expressing 5-HT_2C_R-cMyc and HA-mGluR2 (Figure 2C), or cMyc-5-HT_2A_R and HA-mGluR3 (Figure 2D), and was absent in cells individually transfected with cMyc-5-HT_2A_R or HA-mGluR2 and mixed post-transfection (Figures 2C, 2D).

To explore whether this interaction between transcripts relied solely on the interaction between their corresponding proteins, we impeded the translation of the HA-mGluR2 construct by substituting the start codon (ATG) with a stop codon (TAA) in mGluR2 cDNA (TAA-HA-mGluR2) (Figure 2E). As expected, although the *mGluR2* gene could produce a stable transcript (Figure 2D), it was unable to translate a functional protein compared to its wild-type counterpart (Figure 2F). Interestingly, in cells co-transfected with cMyc-5-HT_2A_R and TAA-HA-mGluR2, immunoprecipitation of 5-HT_2A_R by the anti-cMyc antibody still effectively pulled down both *5-HT_2A_R* and *TAA-mGluR2* transcripts (Figure 2D). Thus, the transcripts associate even when the nascent mGluR2 and 5-HT_2A_R polypeptides do not interact, suggesting that an interaction at the mRNA level integrates the transcripts that form the 5-HT_2A_R-mGluR2 heterocomplex.

In addition, such translational-independent association between *5-HT_2A_R* and *mGluR2* was observed to decrease following EDTA treatment (Figure 2G, 2H), which leads to polysome dissociation, resulting in free ribosomal subunits and mRNA (Panasenko et al., 2019). Together, these findings suggest that, while there is no need for the two nascent peptides to interact to achieve the association of mRNAs encoding *5-HT_2A_R* and *mGluR2*, the EDTA-induced detachment of ribosomes from the mRNA strand also reduces the ability of the anti-HA antibody to pull down mRNA transcripts associated with the nascent HA-tagged protein partner.

### Transcriptional association of *5-HT_2A_R* and *mGluR2* in mouse frontal cortex

To investigate the transcript association of *5-HT_2A_R* and *mGluR2* mRNAs within an endogenous system, we conducted RIP assays using frontal cortex samples from male mice. An anti-mGluR2 antibody directed towards the N-terminus was employed to immunoprecipitate the nascent mGluR2 polypeptide, followed by the assessment of associated transcripts through RT-PCR. Our results indicate that in mouse frontal cortex samples, like in HEK293 cells, nascent mGluR2 polypeptides were in complex with *5-HT_2A_R* and *mGluR2* transcripts encoding protomers of the 5-HT_2A_R-mGluR2 heterocomplex (Figure 4I). Specificity of this interaction was confirmed by the absence of RT-PCR signal with *5-HT_2C_R* primers (Figure 4I). Additionally, reverse transcription quantitative PCR (RT-qPCR) assays validated this finding in a separate cohort of animals (Figure 4J, and Figure S3).

**Figure 3.**
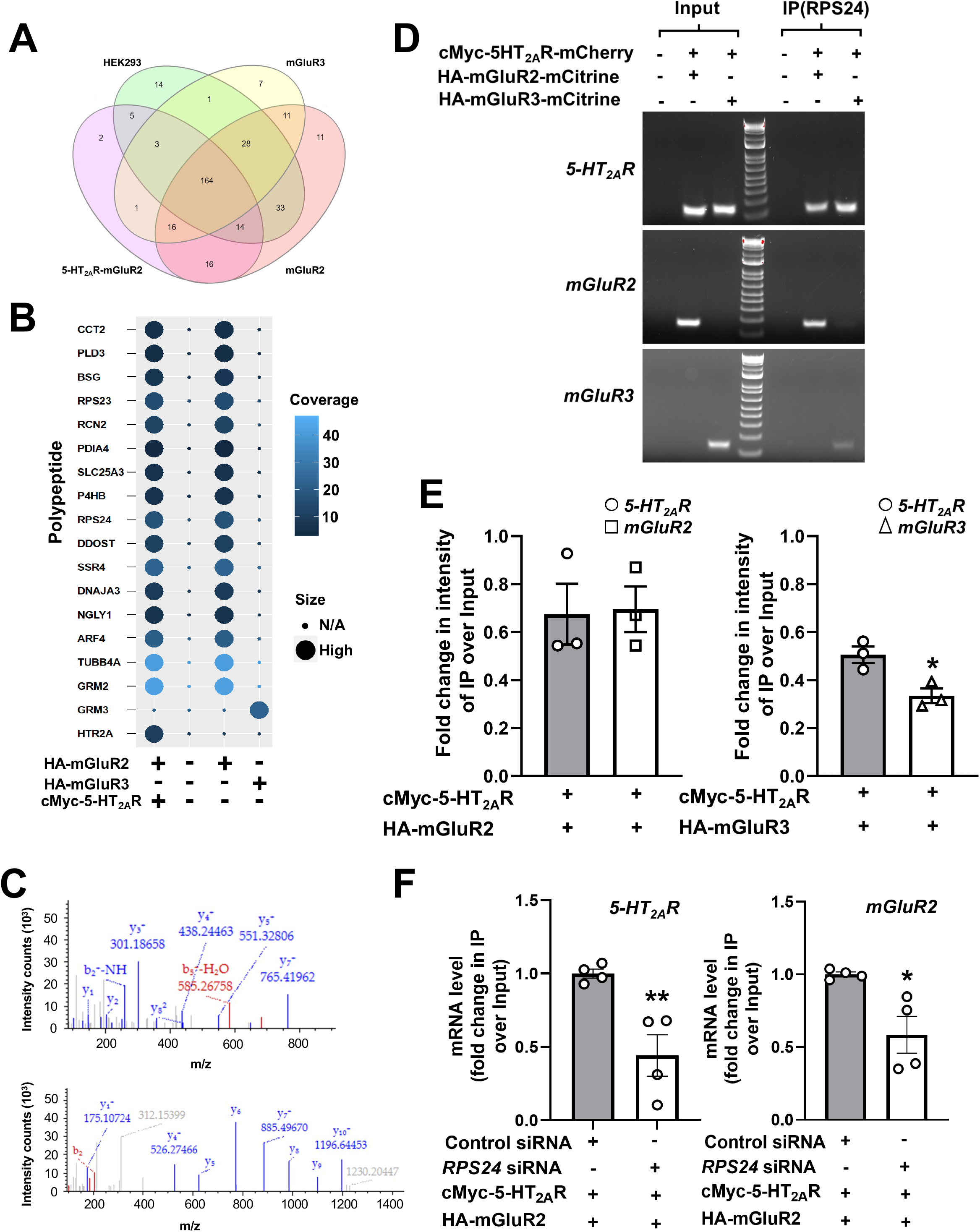
RPS24 facilitates interaction between *5-HT_2A_R* and *mGluR2* throughout translation. (**A**) Venn diagram showing the overlap of identified polypeptides by LC-MS/MS in parental HEK293 cells, cells transfected solely with pcDNA3.1-HA-mGluR2 or pcDNA3.1-HA-mGluR3, and cells co-transfected with pcDNA3.1-HA-mGluR2 and pcDNA3.1-cMyc-5HT_2A_R following RIP assay using the anti-HA antibody. (**B**) Dot plot depicting polypeptides present in HEK293 cells transfected solely with pcDNA3.1-HA-mGluR2 as well as in cells co-transfected with pcDNA3.1-HA-mGluR2 and pcDNA3.1-cMyc-5HT_2A_R, but not in parental HEK293 cells or in cells transfected solely with pcDNA3.1-HA-mGluR3. (**C**) LC-MS/MS analysis identifying with high confidence two tryptic peptides – QMVIDVLHPGK (top) and TTPKVIFVFGFR (bottom) – from RPS24. Mass-to-Charge (m/z) peptide fragments for the y ions (blue) and the b ions (red) are displayed in the spectra **(D, E)** HEK293 cells were co-transfected with pcDNA3.1-cMyc-5-HT_2A_R, and pcDNA3.1-HA-mGluR2 or pcDNA3.1-HA-mGluR3. Images show representative RT-PCR products for *5-HT_2A_R*, *mGluR2* and *mGluR3* transcripts from HEK293 cells before (Input) and after immuno-precipitation (IP) using an anti-RPS24 antibody. Representative image (**D**), and quantification of IP/input band intensities (n = 3 independent experiments) (**E**). (**F**) HEK293 cells were transfected with non-targeting siRNA or *RPS24* siRNA. Forty-eight hours after siRNA transfection, cells were transfected with pcDNA3.1-c-Myc-5-HT_2A_R and pcDNA3.1-HA-mGluR2 constructs. RIP assays were carried out 24 h following DNA transfection using an anti-HA antibody. Subsequently, the RNP complexes underwent processing for RNA isolation and RT-qPCR assays for the detection of *5-HT_2A_R* and *mGluR2* transcripts. Data are shown as fold change of IP/Input (n = 4 independent samples). Unpaired two-tailed Student’s *t*-test (*p < 0.05, ***p < 0.001).

**Figure 4.**
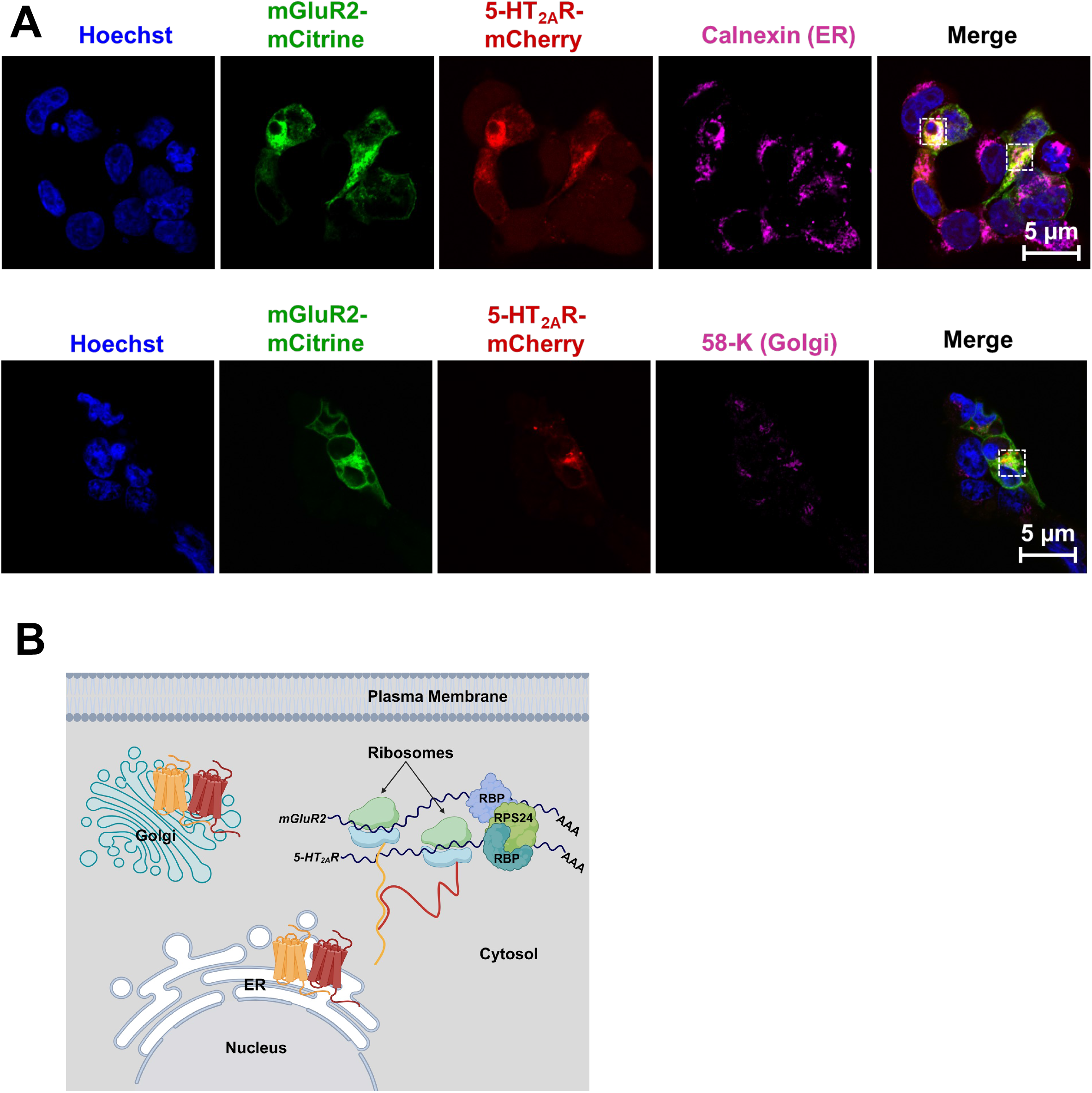
Co-localization of 5-HT_2A_R and mGluR2 along the maturation pathway. (**A**) Confocal micrographs of HEK293 cells transiently transfected with pcDNA3.1-cMyc-5-HT_2A_R-mCherry (red) and HA-mGluR2-mCitrine (green), and stained with anti-calnexin (upper panel, magenta) or anti-58-K (lower panel, magenta) and secondary antibodies, and imaged to detect mCherry, mCitrine, anti-calnexin, and anti-58-K. Nuclei were stained in blue with Hoechst. Scale bars, 5 µM. (**B**) Schematic representation of a working model for the mechanism of *5-HT_2A_R* and *mGluR2* transcript association and subcellular localization of the 5-HT_2A_R-mGluR2 heterocomplex during maturation. Following co-translational association within RNP complexes containing RPS24, 5-HT_2A_R and mGluR2 are trafficked to the ER as part of a protein complex. Subsequently, this GPCR heterocomplex is translocated to the Golgi apparatus and directed to the cell membrane or other subcellular compartments.

### Identification of RNA-binding proteins that enable co-translation of 5-HT_2A_R and mGluR2

To explore the potential involvement of RNA-binding proteins (RBPs) in facilitating the co-translation of 5-HT_2A_R and mGluR2 proteins, we conducted mass spectrometry analysis on cells transfected solely with HA-mGluR2 or HA-mGluR3, along with those co-transfected with HA-mGluR2 and cMyc-5HT_2A_R, comparing them to parental HEK293 cells. As above, following the isolation of RNP complexes through a conventional RIP assay using an anti-HA antibody, the immunoprecipitated fractions underwent LC-MS/MS analysis. This analysis facilitated the identification of 326 polypeptides, which were found to be differentially abundant or absent throughout all four experimental conditions (Figure 3A, and Table S1). Out of these, 16 polypeptides were found in cells that expressed HA-mGluR2 alone, and in cells that co-expressed cMyc-5-HT_2A_R and HA-mGluR2. However, these polypeptides were not detected in cells expressing HA-mGluR3 alone, or in the parental HEK293 cells (Figures 3A, 3B). Interestingly, two of the polypeptides were identified as RBP – RPS23 and RPS24. Further bioinformatics analysis indicated that RPS24 exhibited a higher percentage of coverage, and was identified with high confidence by sequencing of two unique tryptic peptides (QMVIDVLHPGK and TTPKVIFVFGFR) from RPS24 using LC-MS/MS analysis, as shown by the corresponding spectra (Figure 3C). The presence of RPS24 immunoreactivity in parental HEK293 cells and in cells co-transfected with cMyc-5-HT_2A_R and HA-mGluR2 or HA-mGluR3 was further confirmed through immunoblot assays (Figure S4A).

To elucidate the potential involvement of RPS24 in facilitating the interaction between *5-HT_2A_R* and *mGluR2* transcripts, parental HEK293 cells and cells co-transfected with cMyc-5-HT_2A_R and either HA-GluR2 or HA-mGluR3 were subjected to RIP assays employing an anti-RPS24 antibody. Subsequently, the isolated RNP complexes underwent processing for RNA extraction and RT-PCR assays. Interestingly, the transcripts of *5-HT_2A_R* and *mGluR2* were more prominently detected in RNP complexes immunoprecipitated by anti-RPS24, as opposed to *mGluR3* transcripts (Figure 3D). Quantitative analysis of the RT-PCR band intensities revealed significant differences in *mGluR3* intensity compared to *mGluR2* or *5-HT_2A_R* (Figure 3E). Additionally, siRNA-mediated knockdown of *RPS24* in HEK293 cells (Figure S4B) led to a significant reduction in the ability of the anti-HA antibody to pull down HA-mGluR2-associated mRNA transcripts, as determined by RT-qPCR assays (Figure 3F). This further underscores the crucial role of RPS24 in the co-translational assembly of *5-HT_2A_R* and *mGluR2* transcripts.

### Co-localization of 5-HT_2A_R and mGluR2 in the maturation pathway

The association of *5-HT_2A_R* and *mGluR2* transcripts preceding the interaction of their fully functional protein counterparts prompted an investigation into the sub-cellular localization along the maturation pathway immediately following their co-translational assembly on cytosolic RBP-RNA complexes. To visualize the components of the 5-HT_2A_R-mGluR2 heterocomplex within the endoplasmic reticulum (ER) and Golgi apparatus, HEK293 cells co-transfected with cMyc-5HT_2A_R-mCherry and HA-mGluR2-mCitrine were subjected to immunostaining using anti-Calnexin and anti-58K antibodies, respectively. Confocal microscopy analysis revealed a co-localization of cMyc-5HT_2A_R-mCherry and HA-mGluR2-mCitrine fluorescence signal with both ER and Golgi markers (Figure 4A).

Together, these results suggest that RPS24 promotes the assembly of *5-HT_2A_R* and *mGluR2* mRNA transcripts without being contingent on their mRNA-to-protein translation; however, it remains unknown whether RPS24 engages directly with these mRNA transcripts, or alternatively if RPS24 functions as an integral component of a multi-protein complex that facilitates the confluence of these transcripts during the process of translation (Figure 4B).

## DISCUSSION

In this study, we have elucidated the pivotal role of 5-HT_2A_R in governing the transcription and translation processes of mGluR2. Suppression of *5-HT_2A_R* expression using siRNA in HEK293 cells transiently transfected with 5-HT_2A_R and mGluR2 yielded a significant inhibition in the expression of *mGluR2* and its corresponding protein levels, a cross-regulatory event that was not observed in cells co-transfected with 5-HT_2A_R and mGluR3. The observed concurrent decrease in one transcript following the siRNA-induced downregulation of the other implied a potential physical association between the two transcripts. This physical association was further substantiated by RIP assays isolating RNP complexes. Thus, immunoprecipitation of either 5-HT_2A_R or mGluR2, using their tag-specific antibodies, resulted in detection of both transcripts associated with RBPs within RNP complexes containing RPS24 as a pivotal contributor to this molecular assembly. The specificity of this interaction was confirmed by the absence of co-translational association in cells co-transfected with 5-HT_2C_R and mGluR2, or 5-HT_2A_R and mGluR3. To address concerns regarding potential experimental artifacts arising from overexpression, and to evaluate potential co-translational association in a native tissue system, this finding was subsequently validated in mouse frontal cortex tissue samples, where mGluR2 immunoprecipitation in isolated RNP complexes successfully pulled down both *mGluR2* and *5-HT_2A_R* transcripts, but not *5-HT_2C_R*.

RNP complexes represent macromolecular entities comprising RBPs and their target RNAs. The reciprocal regulation of RNAs and RBPs occurs within this intricate complex, where RBPs contribute to RNA processes such as localization, stability, processing, modification and translation. Conversely, RNAs can modulate RBP functions, localization, interactions and stability (Thelen and Kye, 2019; Kelaini et al., 2021). Considering the co-translational association process observed among the *5-HT_2A_R* and *mGluR2* transcripts, one of our principal objectives was focused on the identification of potential RBPs that may be responsible for this mRNA association. Our mass spectrometry analysis identified two ribosomal RBPs – namely RPS23 and RPS24 – in cells co-transfected with *5-HT_2A_R* and *mGluR2*, or mGluR2 alone; but not in cells mock-transfected or transfected with mGluR3 alone. Based on a higher protein sequence coverage, we focused our efforts on RPS24, and, using an anti-RPS24 antibody, we were able to immunoprecipiate from RNP preparations transcripts encoding both *5-HT_2A_R* and *mGluR2*, as well as a scarcer fraction of m*GluR3* mRNA, which suggests that the *5-HT_2A_R* and *mGluR2* mRNA transcripts form part of the same RNP multimer. In shaping the structure of RNA molecules, both intramolecular and intermolecular RNA-RNA interactions (RRIs) assume pivotal roles. These interactions occur either intramolecularly within a single RNA molecule or intermolecularly through engagements among distinct RNA molecules (Van Treeck and Parker, 2018), and can be mediated via either base pairing or RBP intermediates (Xue, 2022). Additionally, the interactions of RBPs with RNA can range from individual protein-RNA element interactions to the intricate assembly of multiple RBPs and RNA molecules, exemplified by the spliceosome (Corley et al., 2020). Several neurogenetic disorders, such as Fragile X syndrome, Spinal muscular atrophy, and Paraneoplastic syndromes arise as consequence of mutations in RBPs (Lukong et al., 2008; Gebauer et al., 2021). The RNA binding domains within RBPs serve as the functional component responsible for RNA binding. Among the numerous RNA binding domains elucidated to date, RNA Recognition Motifs (RRMs) stand out as the most prevalent and extensively investigated RNA binding domain. While the primary role of the *RPS24* gene is to provide instructions for synthesizing various ribosomal proteins, recent studies also suggest that RPS24 may have alternative cytoplasmic functions, such as participating in chemical signaling pathways, regulating cell division and controlling apoptosis. As one example, siRNA-mediated knockdown of *RPS24* resulted in deficits in the biosynthesis of the 40S subunit of the ribosomes, the subunit that binds RNA and mediates translation (Choesmel et al., 2008).

Further investigation will be necessary to determine whether RPS24 directly binds the *5-HT_2A_R* and *mGluR2* mRNA transcripts, or alternatively, if RPS24 is a part of a multi-protein complex that facilitates the association of these transcripts during translation. Additional investigation will also be essential to elucidate the specific region of the mRNAs recognized by RPS24 and/or other RBPs that enable the inclusion of the two coding sequences responsible for the translation of *5-HT_2A_R* and *mGluR2* within the same RNA-protein complex. Moreover, our data do not exclude the possibility that RPS24 influences *mGluR3* transcription through alternative pathways.

An important finding is the lack of effect of *mGluR2* siRNA on *5-HT_2A_R* expression whereas *5-HT_2A_R* siRNA leads to a reduction in both *mGluR2* transcription and mGluR2 translation in cells co-expressing these two neurotransmitter GPCRs. These *in vitro* findings replicate previous transcriptional and epigenetic observations in mouse frontal cortex samples; as *mGluR2* mRNA expression and mGluR2 density assessed by RT-qPCR and radioligand binding assays with the mGluR2/3 antagonist [^3^H]LY341495, respectively, were downregulated in the frontal cortex *5-HT_2A_R-KO* mice (Kurita et al., 2013); whereas 5-HT_2A_R expression (mRNA and binding with the 5-HT_2A_R antagonist [^3^H]ketanserin) was unaffected in the same cortical region of *mGluR2-KO* animals (Moreno et al., 2011). A potential mechanism underlying this unidirectional rather than bidirectional crosstalk in transcriptional regulation may be related to epigenetic processes that involve histone deacetylase 2 (HDAC2) (Graff and Tsai, 2013), as it was reported that chronic administration of the 5-HT_2A_R pharmacological blocker and antipsychotic medication clozapine reduces expression and density of mGluR2 through a process that involves an increase in the recruitment of HDAC2 at the *mGluR2* promoter in mouse frontal cortex; an effect that occurs in concert with repressive epigenetic modifications in histone tails including a decrease histone H3 acetylation (H3ac) (Kurita et al., 2012; de la Fuente Revenga et al., 2019), which correlates with transcriptional activation, and an increase in histone H3 trimethylation at lysine 27 (H3K27me3), a repressive epigenetic modification marker (Bastle and Maze, 2019). While we have demonstrated translation-independent association of mRNAs encoding *5-HT_2A_R* and *mGluR2*, further work needs to be completed to clarify the mechanisms by which this unidirectional transcriptional crosstalk occurs in both *in vitro* and *in vivo* models.

We also observed that 5-HT_2A_R and mGluR2 co-localize with markers of intracellular components of the maturation pathway, including the ER and Golgi apparatus. Considering our previous data revealing that the effect of 5-HT_2A_R on localization of mGluR2 within endosomal compartments required GPCR heteromerization (Toneatti et al., 2020), it will be interesting to expand this analysis in further studies to evaluate physical proximity between 5-HT_2A_R and mGluR2 within the intracellular components of maturation pathway. For the most part, GPCRs are primarily localized at the cell surface, especially in cells not exposed to an agonist (Ferguson, 2001). However, certain GPCRs, including the 5-HT_2A_R (Lopez-Gimenez et al., 2008; Magalhaes et al., 2010; Toneatti et al., 2020; Vargas et al., 2023), have a significant intracellular presence. Signal sequences are crucial in the initial stages of intracellular transport of GPCRs. They participate in targeting nascent chains to the ER membrane, initiating the integration of newly synthesized peptides into this compartment (Rutz et al., 2015). Intriguingly, unlike its counterpart the 5-HT_2C_R (Jahnsen and Uhlen, 2012), the N-terminal of the 5-HT_2A_R lacks a putative cleavable signal peptide. Further research will be necessary to better understand the potential role of this mRNA association in the biosynthesis and maturation of the 5-HT_2A_R-mGluR2 heterocomplex (Bulenger et al., 2005).

In conclusion, our collective data indicate an association between mRNA transcripts encoding 5-HT_2A_R and mGluR2 in both mammalian cells and mouse frontal cortex, along with their corresponding nascent polypeptides. This association is primarily facilitated by a complex of RBPs, with RPS24 among the key contributors. To the best of our knowledge, this represents the first example of translation-independent association of *GPCR* mRNAs, and these findings may provide a new route to modulate neural transcriptional plasticity processes.

## ACKNOWLEDGMENTS

We thank Carlos Escalante, Joseph Landry and Diomedes Logothetis for their critical review of the manuscript, and Mikhail Dozmorov for his help with the Seurat’s dot plot tool. Proteomics expertise and LC-MS/MS analysis was provided by Adam M. Hawkridge, Charles Lyons and Arjun Rijal at the VCU Massey Comprehensive Cancer Center Proteomics Shared Resource whereas Microscopic imaging expertise was provided by Tytus Bernas at the VCU Massey Comprehensive Cancer Center Microscopy Shared Resource, both supported, in part, with funding from NIH-NCI Cancer Center Support Grant P30 CA016059. This work was supported by NIH R01MH084894 to J.G.-M. Schematic models were created using Biorender.com.

## AUTHOR CONTRIBUTIONS

Conceptualization, S.S. and J.G.-M.; funding acquisition, J.G.-M.; investigation, S.S., supervision, J.G.-M.; writing – original draft, S.S.; writing – review and editing, S.S. and J.G.-M.

## DECLARATION OF INTERESTS

J.G.-M. has as sponsor research contract with Terran Biosciences. S.S. declares no conflict of interest.

## METHODS

### Plasmid construction

The constructs pcDNA3.1-cMyc-5-HT_2A_R-mCherry, pcDNA3.1-5-HT_2C_R-cMyc-mCitrine, pcDNA3.1-HA-mGluR2-mCitrine, and pcDNA3.1-HA-mGluR3-mCitrine have previously been described (Moreno et al., 2016). Introduction of mutation (ATG to TAA) into the pcDNA3.1-HA-mGluR2-mCitrine construct was performed with the QuikChange II Site-Directed Mutagenesis Kit (Agilent, Catalog no. 200523) (for primer pairs, see Table S2). All the constructs were confirmed by DNA sequencing.

### Transient Transfection of HEK293 cells

Human embryonic kidney (HEK293) cells (ATCC: CRL-1573) were maintained in Dulbecco’s modified Eagle’s medium (DMEM) supplemented with 10% dialyzed fetal bovine serum (dFBS) and 1% penicillin/streptomycin (Gibco) at 70-80% confluency in a 5% CO_2_ humidified atmosphere.

### Mouse brain samples

On the day of the experiment, male mice (C57BL/6J) were killed by cervical dislocation, and bilateral frontal cortex samples (bregma 1.90 to 1.40 mm) stored at −80°C until tissue processing. All procedures were conducted in accordance with the National Institutes of Health (NIH) guidelines, and were approved by the Virginia Commonwealth University Animal Care and Use Committee.

### RNA interference

For small interfering RNA (siRNA) assays, cells were transfected with non-targeting siRNA (Santa Cruz Biotechnology, Inc.), *5-HT_2A_R* siRNA, *mGluR2* siRNA or *RPS24* siRNA (for siRNA target sequences, see Table S3) (Sigma) using Oligofectamine (Invitrogen), following manufacturer’s instructions. Forty-eight hours after siRNA transfection, cells were transfected with pcDNA3.1-cMyc-5-HT_2A_R-mCherry, pcDNA3.1-HA-mGluR2-mCitrine and/or pcDNA3.1-HA-mGluR3-mCitrine using Lipofectamine 3000 (Invitrogen). Assays including RNA and protein extraction, and isolation of RNP complexes were carried out 24 h following DNA transfection.

### Ribonucleoprotein complex isolation

RNP complexes were isolated using a RiboCluster Profiler TM RIP-Assay Kit (Medical & Biological Laboratories, catalog no. RN1001), according to manufacturer’s protocol. Briefly, HEK293 cells were transiently co-transfected (1:1 ratio) with pcDNA3.1-cMyc-5-HT_2A_R-mCherry, pcDNA3.1-5-HT_2C_R-cMyc-mCitrine, pcDNA3.1-HA-mGluR2-mCitrine, pcDNA3.1-HA-mGluR3-mCitrine, and or pcDNA3.1-TAA-HA-mGluR2-mCitrine using Lipofectamine 3000. Twenty-four hours post DNA transfection cells were processed for isolation of RNP complexes. For this, immunoprecipitation assays were performed for 3-h at 4°C using anti-cMyc (Cell Signaling technology, catalog no. 2276), anti-HA (Cell Signaling technology, catalog no. 2367), or anti-RPS24 (abcam, Catalog no. ab196652) antibodies. This was followed by RNA isolation, DNase 1 treatment, cDNA preparation and analysis of target RNAs using RT-PCR and RT-qPCR assays. For polysome dissociation assays, 25 mM EDTA (ethylenediaminetetraacetic acid; Sigma) or vehicle was to the RIP lysis buffer during the isolation of RNP complexes.

Mouse frontal cortex samples were homogenized using Qiagen TissueRuptor II (120 V, 60 Hz, and Catalog no. 9002755) in the lysis buffer, and proceeded as recommended by manufacturer’s protocol (see above). Immunoprecipitation of mGluR2 was performed overnight at 4°C using a mouse monoclonal anti-mGluR2 N-terminal antibody (Abcam, Catalog no. ab15672), followed by RNA isolation, DNase 1 treatment, cDNA preparation and analysis of target RNAs using RT-PCR and RT-qPCR assays (for primer pair sequences, see Table S2). Specificity of the anti-mGluR2 antibody has been previously reported using *mGluR2-KO* mice (Moreno et al., 2016).

### RNA isolation, RT-PCR, and RT-qPCR

RT-PCR and RT-qPCR assays were performed as previously reported (Moreno et al., 2011; Kurita et al., 2012), with minor modifications (for primer pair sequences, see Table S2). Briefly, total RNA was isolated using RNeasy Mini Kit (Qiagen, catalog no.74104), according to manufacturer’s protocol. Total RNA (2 µg) was then reverse transcribed using High-Capacity cDNA Reverse Transcription Kit (Applied Biosystems, catalog no. 4374966), according to manufacturer’s instructions.

For RT-PCR assays, cDNA (1:30 dilution) was utilized for 30-cycle three-step PCR in Mastercycler Ep Gradient Auto thermal cycler (Eppendorf) using Q5® High-Fidelity 2*×* Master Mix (NEB) and 200 nM of each primer. After electrophoresis, DNA bands were visualized under UV light after staining the agarose gel with ethidium bromide. Quantification of DNA band signals was performed using ImageLab (Biorad). To determine the fold-change for a given transcript between two experimental conditions (*e.g*., EDTA versus vehicle in Figure 2H, or cMyc-5-HT_2A_R and HA-mGluR2 or HA-mGluR3 in Figure 3E), DNA band intensities were normalized to the intensities of the same transcript in the input band.

For RT-qPCR assays, cDNA (1:400, 1:100, and 1:30 dilutions for cell, mouse Input, and mouse IP samples, respectively) was utilized for 40-cycle three-step PCR using PowerUp™ SYBR™ Green Master Mix (Applied Biosystems, catalog number A25742) in an Applied Biosystems QuantStudio 6 Flex Real-Time PCR machine. Each transcript in each sample was assayed two times, and the median threshold cycle (C_T_) was used to calculate the fold change values. The housekeeping gene *GAPDH* was used as an internal control for normalization. In RT-qPCR experiments using total cDNA samples from HEK293 cells (*e.g*., Figures 1A,1D,1G), the corrected C_T_ (cC_T_) for a given transcript (*e.g*., *mGluR2*) was calculated as cC_T(*mGluR2*)_ = C_T(*mGluR2*)_ *−*_-_C_T(*GAPDH*)_. To determine the fold-change for a given transcript, the mean of 2^cCT^ for that transcript in all the control samples (*e.g*., control siRNA) was determined first (control mean). The fold change for each transcript was then calculated from the cC_T_ as 2^cCT^/control mean. In RT-qPCR experiments using RIP samples in HEK293 cells (*e.g*., Figure 3F) and mouse frontal cortex samples (*e.g*., Figure 2J), the corrected C_T_ (cC_T_) for a given transcript (*e.g.,* immunoprecipitated *mGluR2* and input *mGluR2*) was calculated as cC_T(*mGluR2*)_ = C_T(*mGluR2*)_ *−* C_T(Input *GAPDH*)_. After this, 2^cCT^ (immunoprecipitated) values were normalized to 2^cCT^ (input) values of the same transcript. To determine the fold-change, the mean of IP/input in all the control samples (*e.g*., control siRNA) was determined first (control mean). The fold change was then calculated for all the experimental conditions as (IP/input)/control mean.

### Immunoblot

Whole cell lysates (Figures 1C, 1F, 1I, 2F, and Figures S1C, S1F, S2C, S2D) were prepared by disrupting cells using 1*×* lysis buffer followed by sonication. For RIP preparations (Figures S2A, S2B, S4A) cells were harvested and lysed using RIP lysis buffer (see above) for 10 minutes on ice, followed by centrifugation at 12000*×*g for 5 minutes at 4°C. The resulting supernatant was mixed with 1*×* Laemmli sample buffer and boiled at 95°C for 5 minutes. For nuclear fraction preparations (Figures S2A, S2B), HEK293 cells were harvested in cold 1*×*DPBS and homogenized in Tris-HCl (50mM, pH 7.4). The homogenate was centrifuged at 1000*×*g for 5 min at 4°C. The resulting supernatant was discarded and the nuclear pellet was re-suspended in 1*×* Laemmli sample buffer, sonicated and boiled at 95°C for 5 minutes. Equal amounts of protein were separated on an 8% SDS-polyacrylamide gel and transferred to a nitrocellulose blotting membrane. After this, membranes were blocked with 0.1%TBS-T containing 2.5% NFDM and 0.5% BSA, and probed overnight at 4°C with primary antibodies: anti-cMyc (Cell Signaling technology, catalog no. 2276, 1:1000), anti-HA (Cell Signaling technology, catalog no. 2367, 1:1000), anti-GAPDH (Cell Signaling technology, catalog no. 2118, 1:1000), anti-Lamin A/C (Cell Signaling technology, catalog no. 4777, 1:2000), anti-*α*-tubulin (Cell Signaling technology, catalog no. 2125, 1:2000) and anti-RPS24 (Abcam, catalog no. ab196652, 1:1000). Blots were then probed with secondary antibodies (Rabbit IgG HRP Linked Whole Ab, Cytiva NA934-1ML and Mouse IgG HRP Linked Whole Ab, Cytiva NA931-1ML), and immunoreactivity detected with the enhanced chemiluminescence system according to the manufacturer’s instructions. These data compared well with the expected molecular weights of cMyc-5-HT_2A_R-mCherry monomer (*∼*81 kDa), dimer (*∼*162 kDa) and oligomers (>250 kDa), 5-HT_2C_R-cMyc-mCitrine monomer (*∼*79 kDa), dimer (*∼*158 kDa) and oligomers (>250 kDa), HA-mGluR2-mCitrine monomer (*∼*123 kDa), dimer (*∼*246 kDa), HA-mGluR3-mCitrine monomer (*∼*126 kDa), dimer (*∼*252 kDa), GAPDH (*∼*36 kDa), anti-Lamin A/C (*∼*65 kDa and *∼*70 kDa), *α*-tubulin (*∼*50 kDa), and RPS24 (∼15 kDa). Fold-change in immunoreactivity was calculated from experimental target and loading control in each lane using ImageLab software (Biorad). Data represent average of fold-change values from 3 independent experiments.

### LC-MS/MS analysis

HEK293 cells were transiently transfected with pcDNA3.1-cMyc-5-HT_2A_R-mCherry, pcDNA3.1-HA-mGluR2-mCitrine and/or pcDNA3.1-HA-mGluR3-mCitrine, or untransfected (mock). Forty-eight hours after post-transfection, cells were processed for RNP complexes isolation and immunoprecipitation using the anti-HA antibody (see above), and then digested using commercially available Preomics iST sample clean-up protocol. Samples containing approximately 1µg-50µg of protein were mixed with 50µl of lysis buffer, followed by an incubation for 10 minutes at 95°C. Samples were then spun at 1000 rpm to eliminate any foam created during sonication. After this, 50µl of DIGEST solution was added to the mixture, incubated at 37°C for 3 hours and centrifuged at 500 rpm. After digestion, 100µl of STOP solution was added and mixed properly. The digest was then centrifuged at 3800 rpm for 3 mins to ensure complete flow through and washed with 200µl of WASH 1 and 200µl of WASH 2 solution followed by centrifugation after each wash. The cartridge was then placed in a fresh collection tube and 100µl of ELUTE solution was added and centrifuged at 3800 rpm for 3 min to ensure complete flow through. This step was repeated once more to ensure maximum recovery. The elutes were then placed in a vacuum evaporator at 45°C until they dried completely.

LC-MS/MS analysis was performed using a Q-Exactive HF-X (Thermo) tandem mass spectrometer coupled to an Easy nLC 1200 (Thermo) nanoflow UPLC system. The LC-MS/MS system was fitted with an easy spray ion source and an Acclaim PepMap (75μm x 2cm) nanoviper C18 (3μm x 100Å) pre-column in series with an Acclaim PepMap RSLC (75μm x 50cm) C18 2μm bead size (Thermo). The mobile phase consists of Buffer A (0.1% formic acid in water) and Buffer B (80% acetonitrile in water, 0.1% formic acid). The peptides were injected onto the above column assembly and eluted with acetonitrile/0.1% formic acid gradient at a flow rate of 300 nL/min for over 1.6 hours. The nano-spray ion source was operated at 1.9 kV. The digests were analyzed using a data dependent acquisition (DDA) method acquiring a full scan mass spectrum (MS) followed by 15 tandem mass spectra (MS/MS) in the high energy C-trap Dissociation HCD spectra. This mode of analysis produces approximately 50,000 MS/MS spectra of ions ranging in abundance over several orders of magnitude. Not all MS/MS spectra are derived from peptides.

LC-MS/MS datasets data were analyzed in Proteome Discoverer (ver 3.0) using the SEQUEST HT search algorithm and a Custom human and contaminant protein database. Proteins were identified at an FDR <0.01. The m/z (mass-to-charge) ratios were analyzed to identify peptide fragments. Based on this, b ions denote peptide fragments containing the N-terminal of the peptide, whereas y ions correspond to peptide fragments that include the C-terminal. The m/z ratios are presented as the mass of the ion divided by its charge.

### Immunocytochemistry

Approximately 30-h post-transfection, HEK293 cells were fixed with either 4% PFA or ice-cold methanol. Coverslips containing cells were washed thrice with 1x PBS to remove PFA/methanol. Cells were then permeabilized using 0.1% Triton X-100 for 5 min at room temperature and subsequently treated with 5% BSA for 1 hour. Primary antibodies targeting endoplasmic reticulum (Calnexin, Invitrogen, Catalog no. MA3-027, 1:500) or Golgi (58K-9, Abcam, Catalog no. ab27043, 1:100) were incubated overnight at 4°C. After washing thrice with 1x PBS, cells were incubated with goat anti-mouse IgG (H+L) Cross-Adsorbed Secondary Antibody, Alexa Fluor™ 647 (Invitrogen, Catalog no. A-21235, 1:500) for 1-h at room temperature. Cells were washed thrice with 1x PBS and nuclei were stained with Hoechst 33342 Solution (Thermo Scientific, Catalog no. 62249, 1:1000 of 1 mg/ml solution) for 10 minutes at room temperature. After washing twice with 1 *×*PBS, coverslips were mounted on glass slides using ProLong™ Diamond Antifade Mountant (Invitrogen, Catalog no. P36961).

Images of fixed cells were acquired using a Carl Zeiss Axio Observer LSM 710 laser scanning confocal microscope with a Plan-Apochromat 63×/1.40 Oil DIC M27 objective. Hoechst 33342 was excited by a 405-nm blue diode, mCitrine by 488-nm multiline argon laser, mCherry by 561-nm green diode laser, and AF-647 was excited with a 633-nm HeNe laser line. Emission signals were acquired in the same order using the following main beam splitter/dichroic beam splitter (MBS/DBS) and emission filters (EF) sets: InVis 405/Mirror EF, 410 to 483 nm; 488/Mirror EF, 495 to 554 nm; 458/561/633/Mirror EF, 582 to 632 nm (mCherry) and 635-730 nm for AF-647. Pinhole was kept constant at 1 airy unit (AU). After acquisition, images were processed using Fiji software.

### Statistical analysis

Statistical significance was assessed by unpaired Student’s *t* test or two-way ANOVA followed by Bonferroni’s *post-hoc* test. All statistical analyses were performed with GraphPad Prism software, version 10 (Graph-Pad Software, Inc), and comparisons were considered statistically significant if p < 0.05. Data are presented as mean ± s.e.m.

---

**Supplementary Figure S1.**
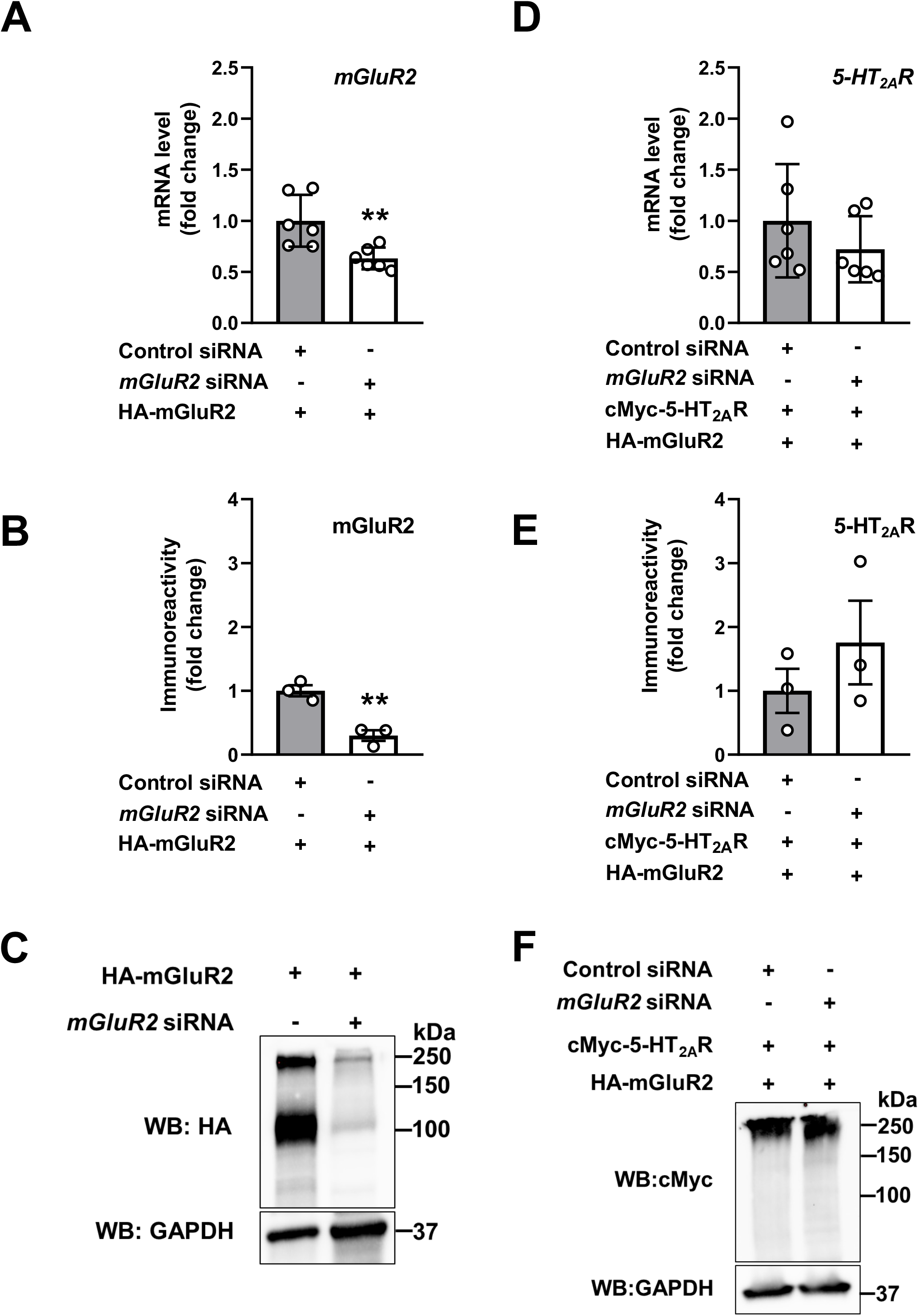
Lack of cross-regulation of 5-HT_2A_R expression by mGluR2. (**A-C**) HEK293 cells were transfected with non-targeting siRNA or *mGluR2* siRNA. Forty-eight hours after siRNA transfection, cells were transfected with pcDNA3.1-HA-mGluR2. RNA and protein extractions were carried out 24 h following DNA transfection. *mGluR2* mRNA was assessed by RT-qPCR (n = 6 independent experiments) (**A**), and mGluR2 immunoreactivity was assessed by Western blot (n = 3 independent experiments) (**B**). Representative immunoblots are shown (**C**). (**D-F**) HEK293 cells were transfected with non-targeting siRNA or *mGluR2* siRNA. Forty-eight hours after siRNA transfection, cells were co-transfected with pcDNA3.1-HA-mGluR2 and pcDNA3.1-c-Myc-5-HT_2A_R. RNA and protein extractions were carried out 24 h following DNA transfection. *5-HT_2A_R* mRNA was assessed by RT-qPCR (n = 6 independent experiments) (**D**), and 5-HT_2A_R immunoreactivity was assessed by Western blot (n = 3 independent experiments) (**E**). Representative immunoblots are shown (**F**). Unpaired two-tailed Student’s *t*-test (p > 0.05).

**Supplementary Figure S2.**
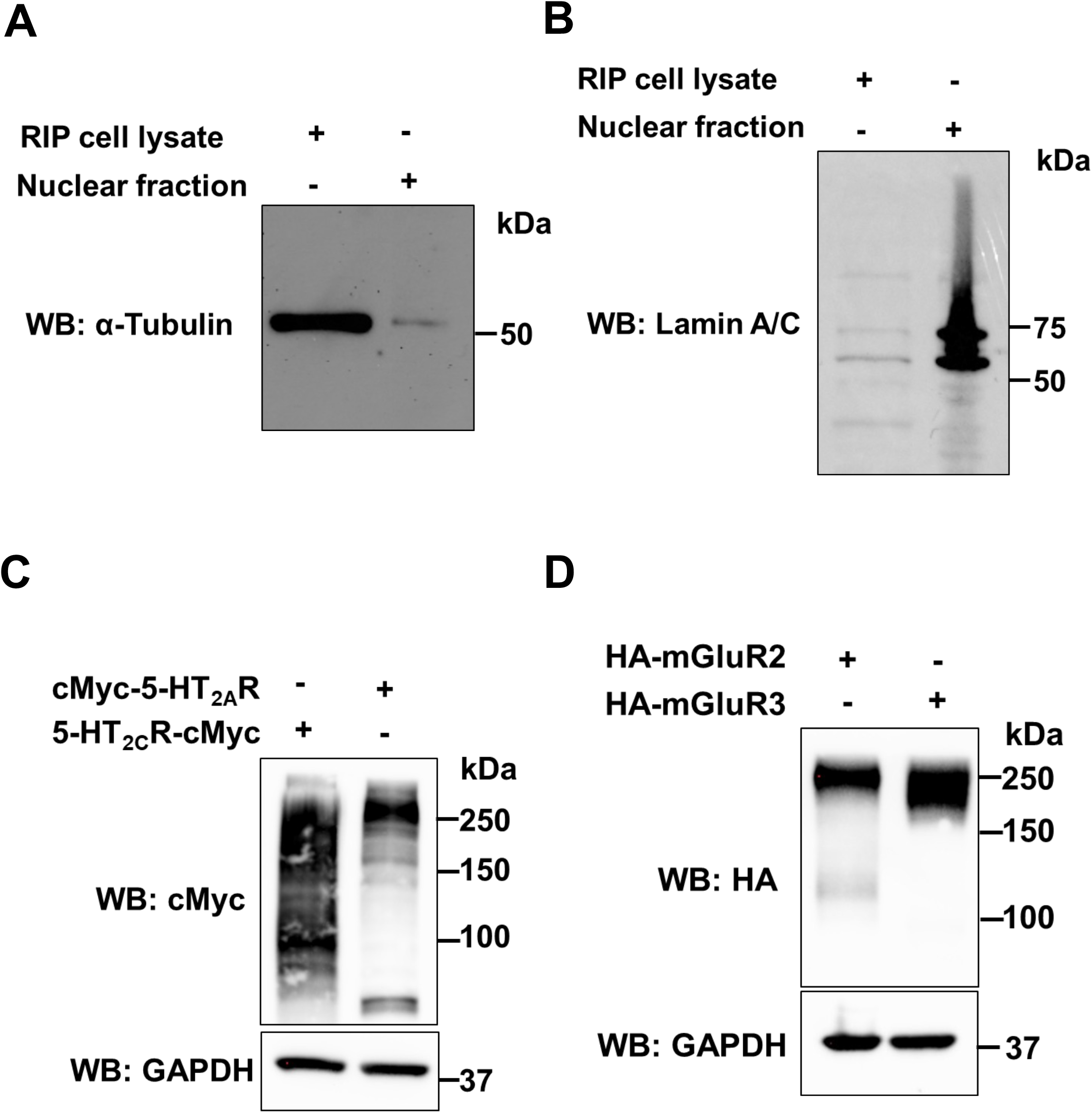
Characterization of 5-HT_2A_R, 5-HT_2C_R, mGluR2 and mGluR3 constructs. **(A, B)** Validation of cytosolic fractionation of RNP complexes using RIP Assay kit by using a cytosolic marker α-Tubulin **(A)** and a nuclear marker Lamin A/C **(B)**. **(C)** Immunoblot with an anti-cMyc antibody in HEK293 cells transiently transfected with pcDNA3.1-c-Myc-5-HT_2A_R or pcDNA3.1-5-HT_2C_R-cMyc. **(D)** Immunoblot with an anti-HA antibody in HEK293 cells transiently transfected with pcDNA3.1-HA-mGluR2 or pcDNA3.1-HA-mGluR3.

**Supplementary Figure S3.**
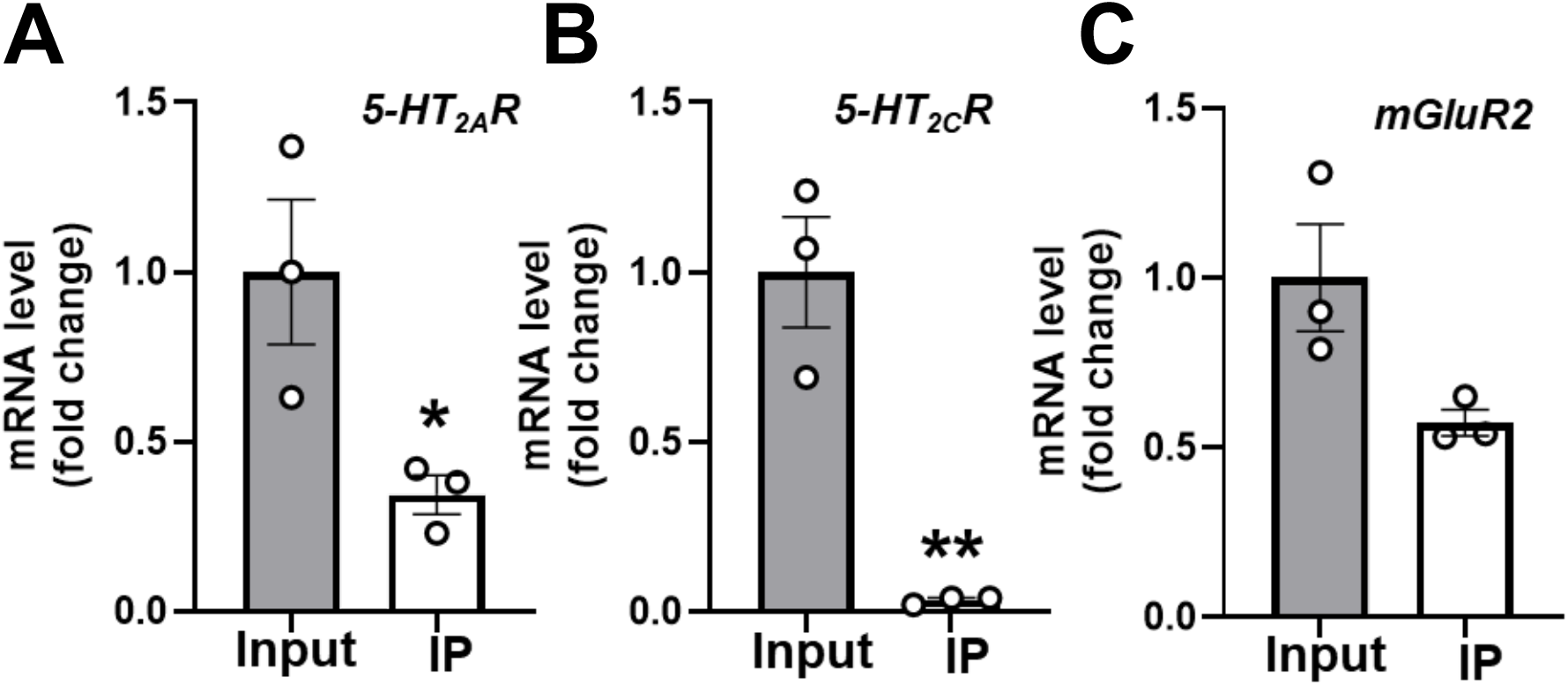
RT-qPCR assays in input and IP samples from mouse frontal cortex. **(A-C)** Mouse frontal cortex samples were subjected to RIP assays employing an anti-mGluR2 antibody. Subsequently, input and IP samples underwent processing for RNA isolation and RT-qPCR assays for the detection of *5-HT_2A_R* **(A)**, *5-HT_2C_R* **(B)** and *mGluR2* **(C)** transcripts (n = 3 mice). Unpaired two-tailed Student’s *t*-test (*p < 0.05, **p < 0.01).

**Supplementary Figure S4.**
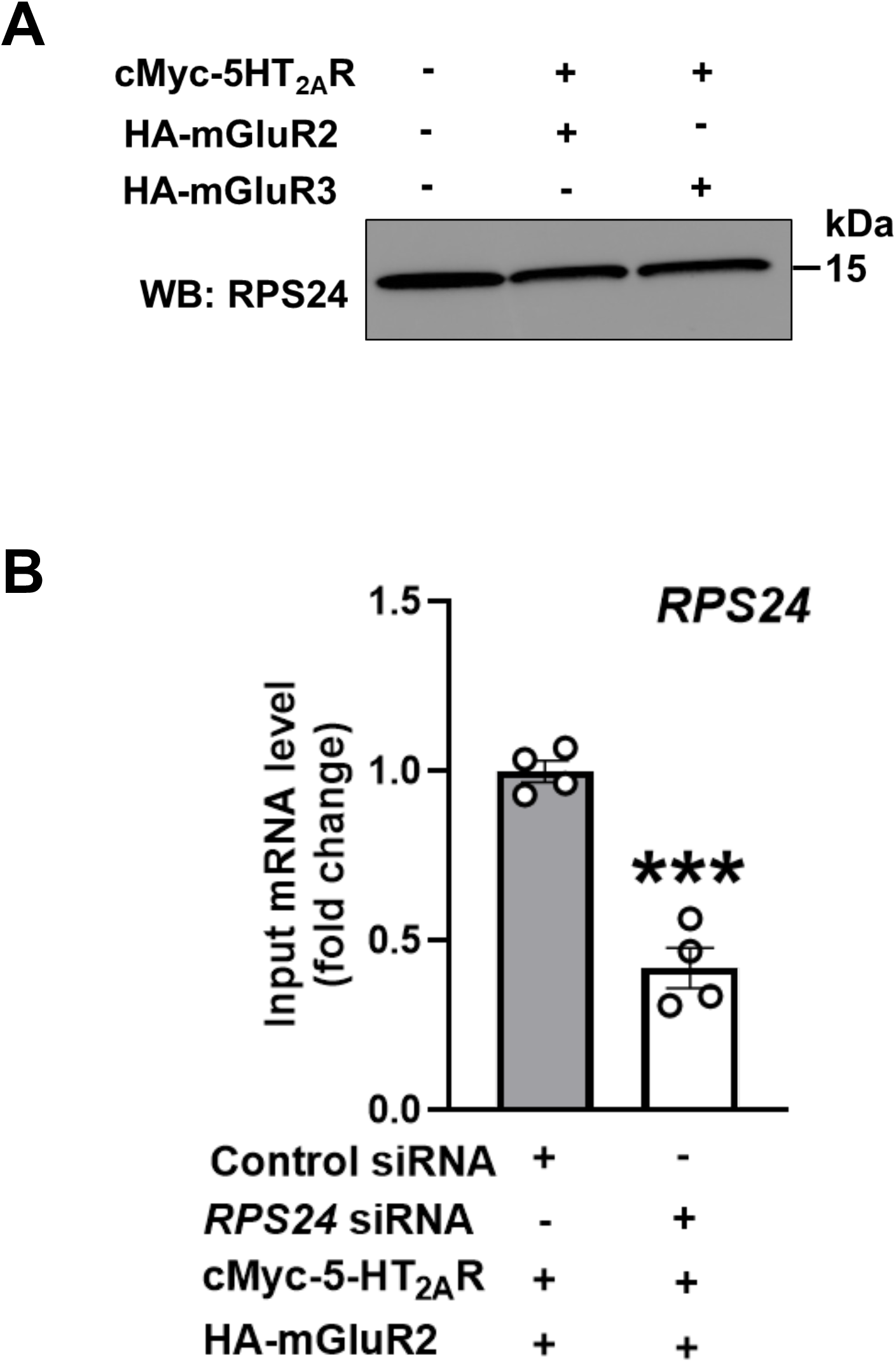
Endogenous expression of RPS24 in HEK293 cells and its siRNA-mediated knockdown in HEK293 cells. **(A)** Immunoblot depicting RPS24 immunoreactivity in RIP preparations from parental HEK293 cells, as well as cells co-transfected with pcDNA3.1-cMyc-5HT_2A_R, and pcDNA3.1-HA-mGluR2 or pcDNA3.1-HA-mGluR3. **(B)** HEK293 cells were transfected with non-targeting siRNA or *RPS24* siRNA. Forty-eight hours after siRNA transfection, cells were co-transfected with pcDNA3.1-cMyc-5HT_2A_R and pcDNA3.1-HA-mGluR2 constructs. RNA extractions were carried out 24 h following DNA transfection in Input samples. *RPS24* mRNA was assessed by RT-qPCR (n = 4 independent samples). Unpaired two-tailed Student’s *t*-test (***p < 0.01).

**Supplementary Table S2:**
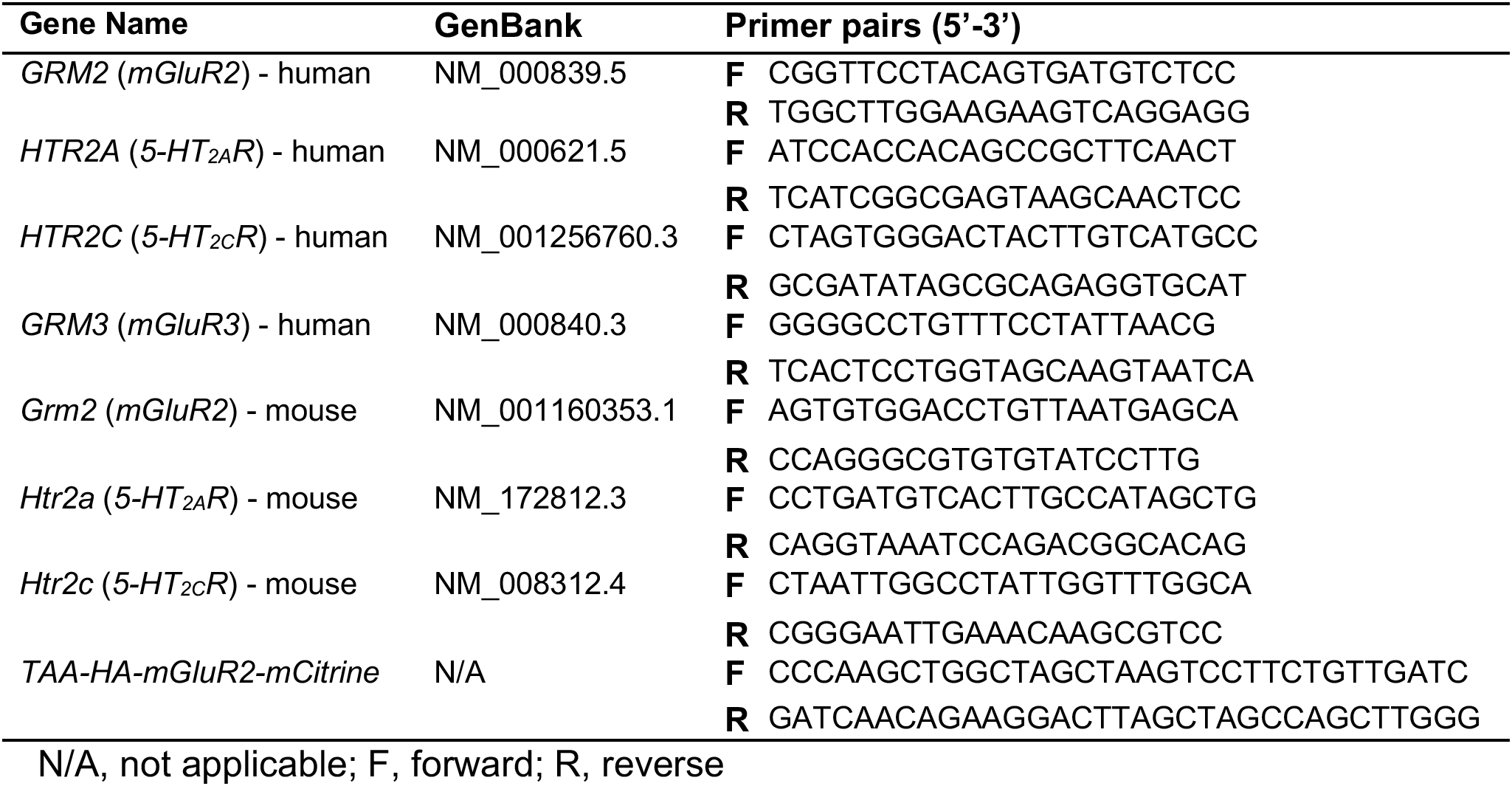
Primer sequences for RT-PCR, RT-qPCR and site-directed mutagenesis.

**Supplementary Table S3:**
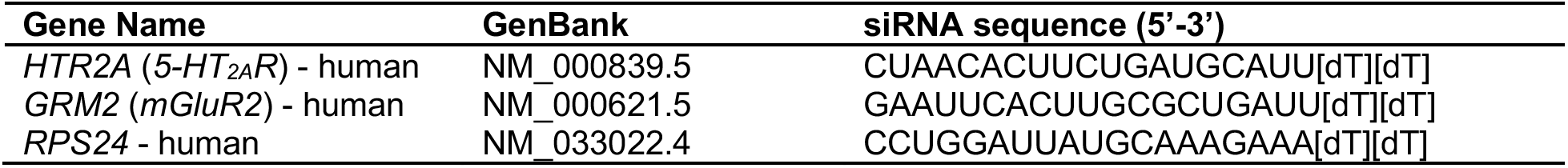
siRNA primer sequences.

